# Three-photon in vivo imaging of neurons and glia in the medial prefrontal cortex with sub-cellular resolution

**DOI:** 10.1101/2024.08.28.610026

**Authors:** Falko Fuhrmann, Felix C. Nebeling, Fabrizio Musacchio, Manuel Mittag, Stefanie Poll, Monika Müller, Eleonora Ambrad Giovannetti, Michael Maibach, Barbara Schaffran, Emily Burnside, Ivy Chi Wai Chan, Alex Simon Lagurin, Nicole Reichenbach, Sanjeev Kaushalya, Hans Fried, Stefan Linden, Gabor C. Petzold, Gaia Tavosanis, Frank Bradke, Martin Fuhrmann

**Author notes:** Corresponding author: Prof. Dr. Martin Fuhrmann Neuroimmunology and Imaging Group German Center for Neurodegenerative Diseases (DZNE) Venusberg-Campus 1/99 53127 Bonn Germany. These authors contributed equally (Falko Fuhrmann, Felix C. Nebeling, Fabrizio Musacchio).

## Abstract

The medial prefrontal cortex (mPFC) is important for higher cognitive functions, including working memory, decision making, and emotional control. *In vivo* recordings of neuronal activity in the mPFC have been achieved via invasive electrical and optical approaches. Here we apply low invasive three-photon *in vivo* imaging in the mPFC of the mouse at unprecedented depth. Specifically, we measure neuronal and astrocytic Ca^2+^-transient parameters in awake head-fixed mice up to a depth of 1700 µm. Furthermore, we longitudinally record dendritic spine density (0.41 ±0.07 µm^-1^) deeper than 1 mm for a week. Using 1650 nm wavelength to excite red fluorescent microglia, we quantify their processes’ motility (58.9 ±2% turnover rate) at previously unreachable depths (1100 µm). We establish three-photon imaging of the mPFC enabling neuronal and glial recordings with subcellular resolution that will pave the way for novel discoveries in this brain region.

## Introduction

The invention and application of two-photon (2P) laser scan microscopy for deep tissue *in vivo* imaging has greatly advanced our understanding about the brain (*1, 2*). It has been applied to visualize and analyze different cell types in various brain regions and model organisms (*3–8*). Indicators, either synthetic or genetically encoded, have been applied to measure spatio-temporal dynamics of structure and function in networks of cells, individual cells or sub-cellular structures (*9–22*). However, light scattering limited the depth penetration of two-photon excitation in brain tissue usually to less than a millimeter with cellular resolution (*23, 24*). Using higher wavelengths above 1300 nm for three-photon (3P) excitation allowed this limit to be exceeded in the brain (*25*). Even functional Ca^2+^-imaging of hippocampal CA1 neurons became possible in young mice (*26*). In addition to neurons, non-neuronal cells, including astrocytes, oligodendrocytes and microglia have been imaged at increased depth in and below the somatosensory cortex (*27, 28*). Advantages of 3P-imaging have also been demonstrated in different model organisms, including zebrafish and *Drosophila* (*29–32*).

A brain region that has not been targeted with 3P-imaging up to now is the medial prefrontal cortex. The mPFC is involved in several higher cognitive functions, including memory, decision making and emotions (*33–35*). Disturbances in the mPFC are hallmarks of different diseases including schizophrenia (*36*), autism spectrum disorders (*37*), and Alzheimer’s disease (*38*). Hence, this region is of high relevance for many neuroscientists. Optical access to record neuronal activity of mPFC neurons can be achieved by two invasive approaches. First, implantation of gradient refractive index (GRIN) lenses in combination with head-held microscopes can be used to record from mPFC neurons (*39–41*). GRIN lenses are available with different diameters ranging from 500 µm up to more than a millimeter (*42*). Cortical regions above the mPFC have to be penetrated and are partially damaged by their implantation. Second, microprisms have been implanted into the fissure between both hemispheres compressing the contralateral hemisphere opposite to the imaged ipsi-lateral hemisphere (*43*). Both methods are restricted by their limited imaging depth (approximately one hundred microns) starting from the implant. Limited imaging depth similarly applies for sub-cellular resolution *in vivo* imaging of dendritic spines in the PFC. Even with invasive microprism implantation only apical dendrites of L5 pyramidal neurons in the mPFC could be resolved with sub-cellular resolution (*44, 45*).

Here we performed three-photon *in vivo* imaging through a cranial window to overcome these limitations. We achieved sub-cellular resolution imaging to measure spine density on the same L5 basal dendrites in the prelimbic area of the mPFC over a week. We recorded neuronal and astrocyte activity via Ca^2+^-imaging throughout the entire dorsal column reaching the infralimbic areas of the mPFC. Time-lapse imaging of td-Tomato expressing microglia and their fine processes at 1650 nm excitation wavelength at a depth of >1000 µm, demonstrated the low invasiveness of our 3P-imaging approach, and revealed the kinetics of microglial fine processes’ motility in the mPFC.

## Results

### Access of the medial prefrontal cortex with 3P-imaging at 1.6 mm depth

We built a 3P-microscope suitable for *in vivo* imaging as shown in **Fig. 1A**. The 3P-microscope consisted of a Thorlabs Bergamo multiphoton setup and a SPIRIT / NOPA laser combination from Spectra Physics tunable between 1300 – 1700 nm (Newport, SPIRIT 1030-70 and NOPA-VISIR+I). The laser repetition rate was maintained at 2 MHz and the microscope was equipped with 900-1900 nm coated optics in the primary scan path to enable transmission of 1300 to 1700 nm excitation light. To validate the capabilities of our 3P-microscope for *in vivo* imaging of the mPFC we chronically implanted rectangular cranial windows above the sagittal sinus to gain optical access to all areas and layers of the mPFC (**Fig. 1B**). Imaging was performed in head-fixed anesthetized YFP-H transgenic mice expressing yellow fluorescent protein (YFP) in a subset of excitatory neurons in different layers of the mPFC (**Fig. 1C**). These experimental conditions enabled us to image up to 1.6 mm deep from the brain surface, covering anterior cingulate, prelimbic and infralimbic areas (**Fig. 1D**; **movie S1)**. To compare 3P-imaging with 2P-imaging, a Ti:sapphire laser (Chameleon Ultra II, Coherent, Santa Clara, USA) was additionally integrated into the excitation light-path. The identical z-stack was imaged sequentially with 1300 nm excitation (3P) and 920 nm excitation (2P) (**Fig. 1E**; **movies S2, S3)**. We achieved imaging two times deeper with 3P-excitation compared to 2P-excitation, using the same maximum laser power (200 mW at maximum depths). The necessary excitation power at 920 nm (2P) increased up to an imaging depth of 300 µm and started to plateau from 400 – 700 µm depth. The necessary excitation power at 1300 nm wavelength (3P) was low up to a depth of 400 µm, then exponentially increased up to a depth of 800 µm and started to plateau at 1000 µm (**Fig. 1F**). The signal to background ratio (SBR) of 2P-imaging was superior to 3P-imaging within the first 400 µm, decreasing to less than two at 500 – 600 µm depth. In contrast, the SBR of 3P-imaging only slightly decreased up to a depth of 1000 µm. Only at depths deeper than 1000 µm, the SBR started to decline (**Fig. 1G**).

**Fig. 1.**
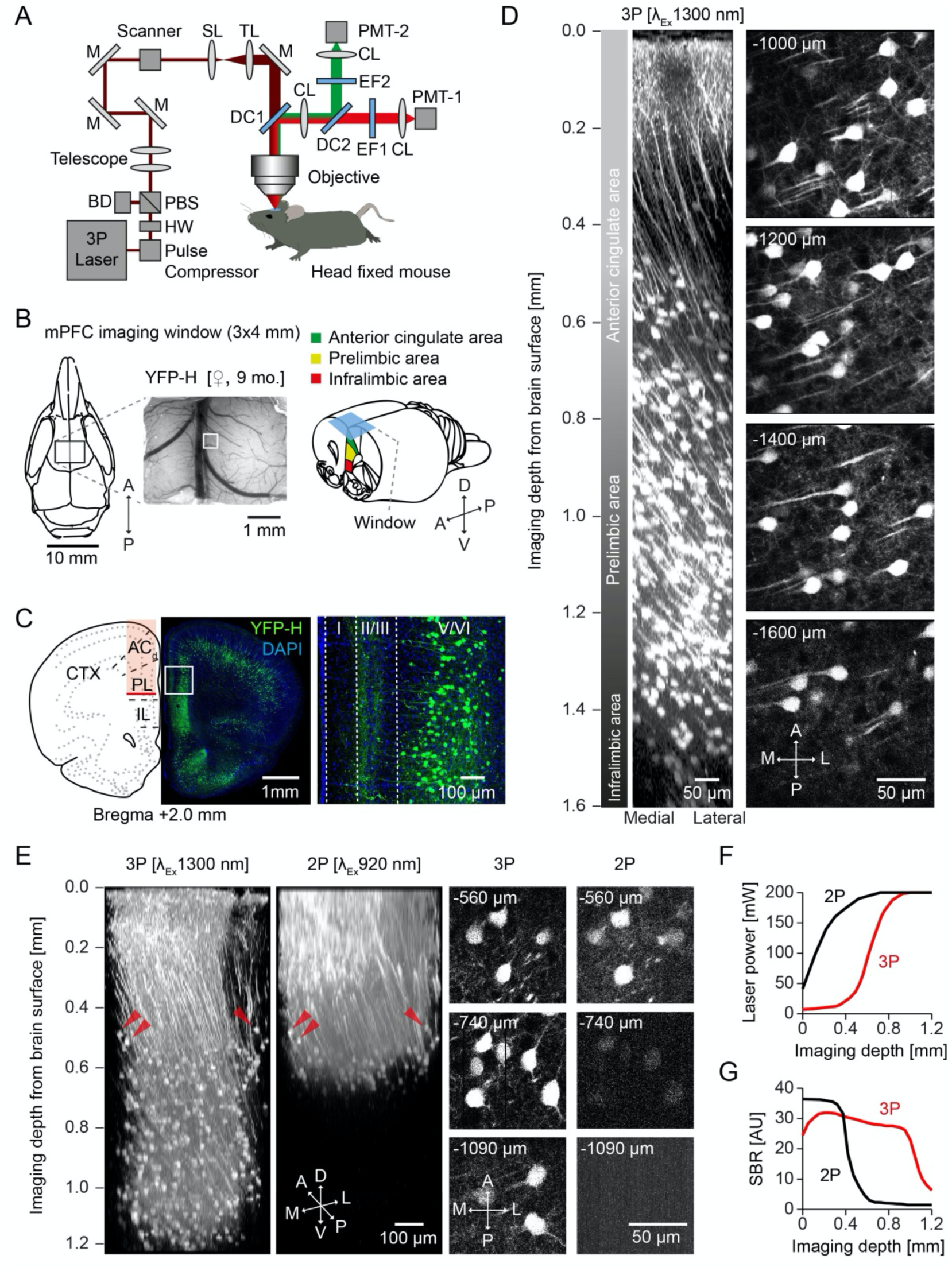
Validation of 3P in vivo imaging of mPFC. **(A)** Setup scheme with light path and optical components. (BD: Beam dump CL: Collection Lens, DC: Dichroic, EF: Emission Filter, HW: Half wave plate, M: Mirror, PBS: Polarizing beam splitter, SL: Scan Lens, TL: Tube Lens) **(B)** mPFC imaging window position on the scull and representative image of the brain surface and vasculature (left, indicated ROI of z-stack shown in D); scheme of the anatomical localization of imaging window in relation to the mPFC subareas (right). **(C)** Confocal image of a coronal section 2 mm anterior of bregma and expression pattern from a YFP-H transgenic mouse in mPFC. Magnification and cellular YFP expression in layer V/VI of mPFC prelimbic area. **(D)** 3D reconstruction of 320 x-y frames from brain surface to 1600µm below taken at a depth increment of 5µm (left) and individual frames at stated imaging depths (right). **(E)** 3D reconstruction and side by side comparison of identical z-stacks recorded with 3P 1300nm and 2P 920nm excitation with individual frames at stated imaging depths. **(F)** Laser Power usage as a function of imaging depth for the z-stacks shown in E. **(G)** Signal to background ratio (SBR) as a function of imaging depth for the z-stacks shown in E.

To achieve high penetration depth with high SBR, we optimized the microscope. Pulse broadening of femtosecond lasers, due to several optics included in the light path, significantly reduces excitation efficiency. Therefore, we included a prism-based compensator from APE (FemtoControl, APE) for dispersion compensation **(Fig. S1A)**. Dispersion compensation was adjusted under visual control of the interferogram measured after the objective. For 1300 nm negative chirp and for 1700 nm, positive chirp was systematically adjusted according to manufacturer’s instructions. To probe its effect on advanced *in vivo* image quality, an identical z-stack was imaged subsequently with pulse compression ON and OFF **(Fig. S1A)**. The comparison of fluorescence histograms at 1 mm depth with or without pulse compression showed more counts of high intensity fluorescence, when pulse compression was switched ON **(Fig. S1B)**. Signal to background ratio was improved by compensation for group delay dispersion throughout all imaging depths **(Fig. S1C)**.

Furthermore we used a 25x water immersion microscope objective with a numerical aperture of1.05 and transmission up to 1700 nm (XLPLN25XWMP2, Olympus) and measured its SBR in comparison to a custom Zeiss 20x, and Nikon 16x – objective, in an identical subsequently recorded cortical volume **(Fig. S1D,E)**. The comparison of fluorescence histograms of the three objectives at 1 mm depths shows highest fluorescent intensities for the Olympus 25x **(Fig. S1D)**. Olympus 25x and Zeiss 20x show similar and enhanced SBR compared to Nikon 16x at all imaging depths **(Fig. S1E)**. Further improvement was achieved by increasing the size of detection optics (size of dichroic, collection lens, BP-filters), which increased the angle to collect more non-ballistic photons in the emission path **(Fig. S1F-I)**. To demonstrate the efficiency, an identical z-stack was recorded in the same mouse increasing the angle from 8° to 14° **(Fig. S1F)**. Highest fluorescence intensities were measured at 1 mm depths with the 14° detector port **(Fig. S1G)**. Highest SBR was achieved with the largest (14°) angle **(Fig. S1H, I)**. Next, we attempted to access the hippocampus with 3P-imaging through the intact cortex **(Fig. S2).** Since it is difficult to penetrate the highly myelinated corpus callosum in adult mice (4 months old), we chose a lateral imaging position to circumvent it **(Fig. S2A).** This enabled us to image hippocampal CA1 neurons deeper than 1 mm. In contrast, with 2P-imaging it was not possible to record YFP-expressing neurons deeper than 700-800 µm **(Fig. S2A-C)**.

Similar to imaging through the corpus callosum, the spinal cord represents an imaging challenge due to optical scattering from myelinated dorsal axons. Therefore, 2P-imaging approaches have been limited to superficial spinal grey matter laminae or ascending dorsal fiber tracts (*46, 47*). We tested our 3P-microscope in adult (3 months old) mouse spinal cord. In a GFP-M mouse, following dorsal laminectomy and dura removal, a spinal cord window was implanted **(Fig. S3A)**. Vertebra were stabilized and the objective placed above grey matter dorsal horn **(Fig. S3B)**. Imaging depths of 390 µm were possible, including axons, cell bodies, dendrites and spines **(Fig. S3C; movie S12)**. Thus, consistent with prior reports, it was not possible to achieve the depths of tissue penetration in spinal cord that we observed in other CNS regions. However, our system allowed for detailed view of neuronal architecture in spinal cord grey matter.

These data demonstrate access to the mPFC using 3P *in vivo* imaging up to a depth of 1600 µm with a chronically implanted cranial window as well as its limitations caused by light scattering myelinated axons.

### Time-lapse 3P-imaging of dendritic spines in the prelimbic area of the mPFC

Recording of subcellular structures, including dendritic spines in deep brain regions such as the prelimbic region of the mPFC can be achieved by implantation of microprisms (*44, 45*). However, the implantation of microprisms is an invasive procedure accompanied by tissue compression or lesions, if incisions or aspirations are necessary for the positioning of the implant. 3P-imaging promises to diminish the invasiveness for subcellular resolution imaging. Therefore, we recorded structural plasticity of dendritic spines in prelimbic areas of the mPFC. We implanted a rectangular cranial window in an aged (16 months old) thy1-GFP-M transgenic mouse (**Fig. 2A**) sparsely expressing green fluorescent protein (GFP) in a subset of excitatory neurons of the cortex (**Fig. 2B**). As in thy1-YFP-H mice, we reached mPFC prelimbic area in thy1-GFP-M transgenic mice with 3P-imaging (**Fig. 2C**; **movie S4)**. Dendritic spines on basal dendrites of layer 5 neurons were clearly visualized at a depth of 900 – 1100 µm (**Fig. 2D**). The measured spine density was 0.41 ±0.07 spines per micrometer (**Fig. 2D**). Of note, we did not use any adaptive optics to correct for wavefront distortions. Furthermore, we repetitively imaged the same dendritic spines over a week visualizing their structural plasticity at a depth of 1000 µm below the brain surface (**Fig. 2E**). These data show that 3P-imaging enables subcellular resolution imaging of structural plasticity of spines over a time-period of one week in the prelimbic area of the mPFC.

**Fig. 2.**
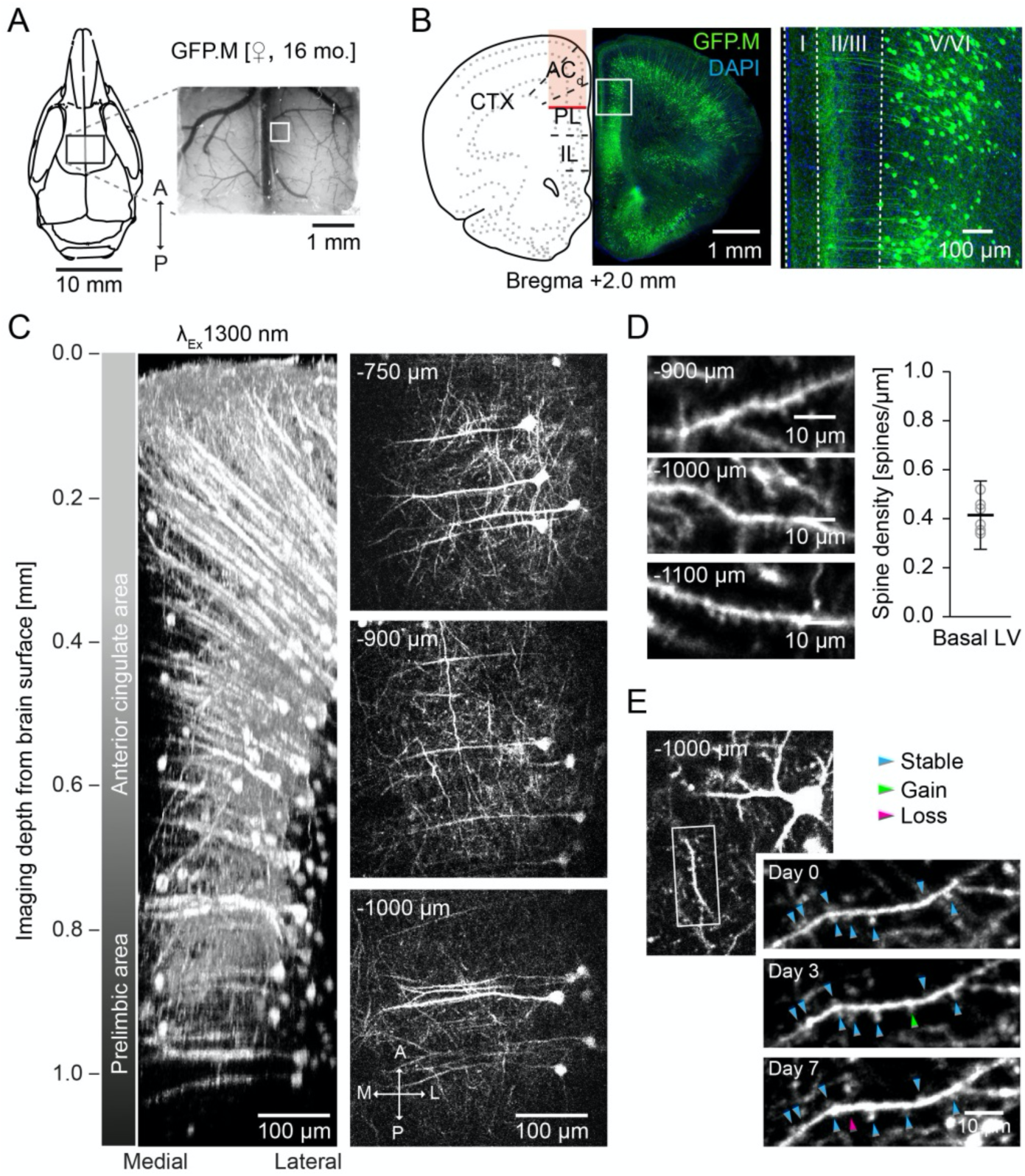
In vivo 3P imaging of dendritic spines in mPFC. **(A)** mPFC window positioning and image of the brain surface and vasculature in a Thy1-GFP-M mouse. **(B)** Schematic and coronal section illustrating the imaging area (left). Zoom of the marked ROI in (B) showing cellular GFP expression in layer V/VI of mPFC prelimbic area (right). **(C)** 3D reconstruction and exemplary images of a z-stack acquired at the ROI marked in (A). **(D)** Exemplary images of three different basal dendrites of L5 neurons with spines in mPFC prelimbic area (left) and the average spine density (n=6, total length=280,5 µm). **(E)** Longitudinal (7d) imaging of dendritic spines on L5 neurons in mPFC prelimbic area. Arrowheads indicate stable (blue), gained (green) and lost (magenta) spines.

### Recording microglial fine processes’ motility in mPFC

Microglia are innate immune cells in the brain. The discovery of motility of their fine processes opened an entire new research field almost 20 years ago (*10, 13*). Up to now, microglia have been imaged in various brain regions, however, the recording depth with 2P-imaging was limited to approximately 300-400 µm. To compare the recording depth between 2P-and 3P-imaging, we implanted cranial windows over the right somatosensory cortex in double transgenic Thy1-YFP-H::Cx3cr1-GFP mice **(Fig. S4A)**. These mice express YFP in a subset of excitatory neurons and GFP in microglia. With 2P*-*imaging at 920 nm microglia and neurons were visualized up to a depth of 600 – 700 µm, whereas 3P-imaging at 1300 nm excitation enabled the recording of microglia in the corpus callosum (800 µm deep) and neurons in dorsal CA1 of the hippocampus **(Fig. S4B, C)**.

Red shifted fluorescent indicators can be excited with less scattering at higher excitation wavelength light and promise improved imaging depth. We used *Cx3cr1-Cre^ERT2^::Rosa26_tdTomato* transgenic mice (*48*) expressing tdTomato upon Tamoxifen-induction in microglia to carry out longitudinal 3P-imaging of microglia in the mPFC with 1650 nm excitation (**Fig. 3A-C**; **movies S5, S6)**. We measured microglial fine process motility (*49*) for a period of 30 min of the very same microglia on two consecutive days (**Fig. 3C-F**). The morphology and position of microglia was unchanged comparing the two time-points (**Fig. 3C**). We measured 44.6 ±4% stable fraction, 27.1 ±4% gained and 28.4 ±3% lost processes on the first imaging time-point. These numbers were comparable to the second time-point 24h later (stable: 41.1 ±2%, gain: 27.4 ±4%, loss: 31.5 ±5%) (**Fig. 3D, E**). Likewise, turnover rate of microglial processes was comparable between the two days (d0: 55.4 ±4%, d1: 58.9 ±2%) (**Fig. 3F**). To assess potential phototoxic or heating effects, we stained for heatshock protein (Hsp40) (**Fig. 3G-I**). While we detected 11.5% Hsp40-positive cells in a mouse model of glioblastoma, we did not find any Hsp40-positive cells in the ipsi-or contra-lateral side of the previously in vivo imaged mPFC region (**Fig. 3I**). Applying 3P-imaging in the mPFC, we repetitively measure the kinetics of microglial motility, underscoring minimal invasiveness of our approach.

**Fig. 3.**
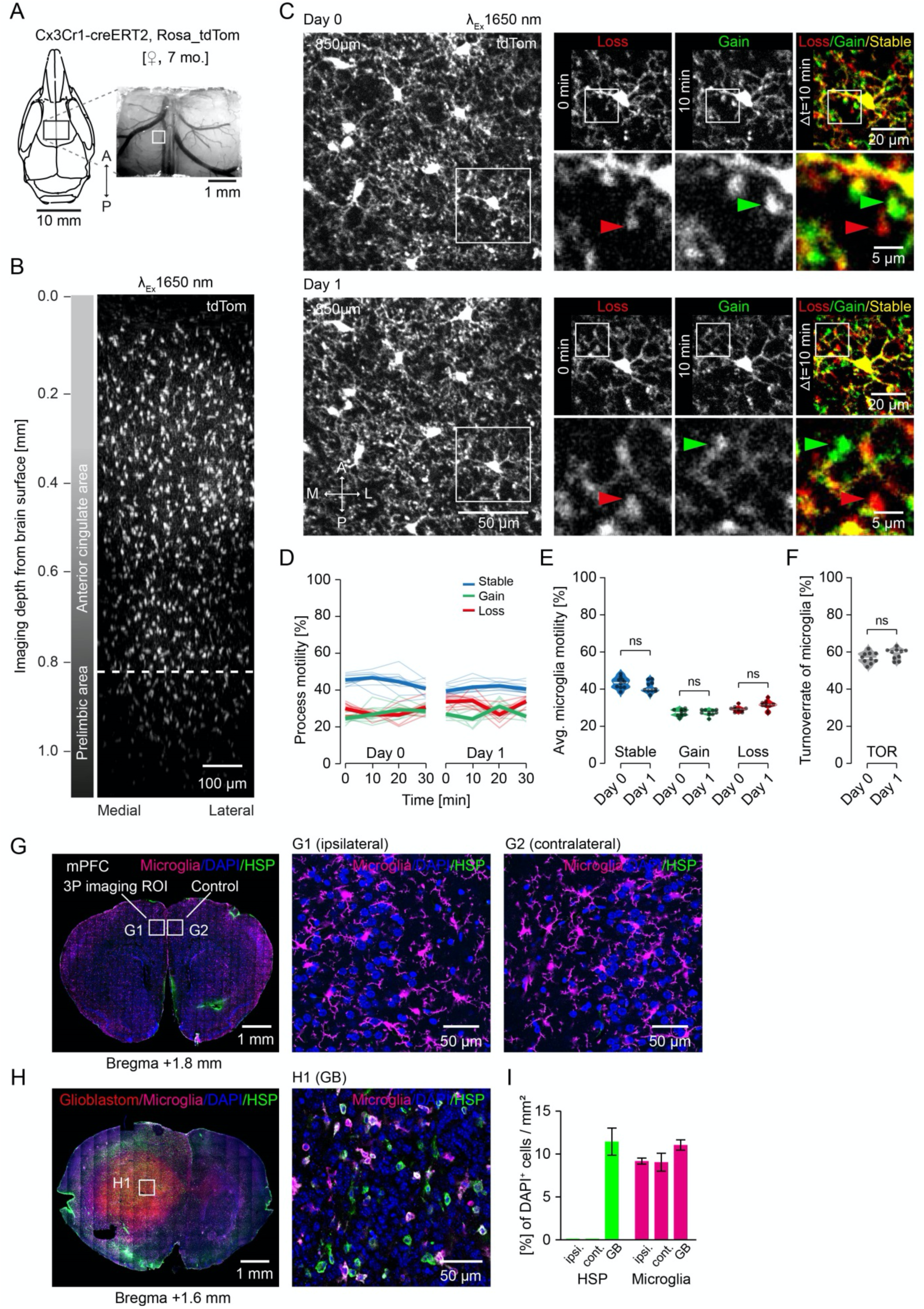
In vivo 3P imaging of microglia in mPFC. **(A)** mPFC window positioning and image of the brain surface and vasculature (indicated ROI of z-stack shown in B) of a 7 months old Cx3Cr1-creER2 Rosa tdTomato mouse. **(B)** 3D reconstruction of 212 x-y frames from brain surface to 1060 µm below taken at a depth increment of 5 µm with 1650 nm Excitation. Dashed line indicating in vivo imaging depth of microglial fine process motility recordings shown in C). **(C)** Average intensity projections of 9 frames (z-spacing: 3µm) 850 µm deep of the very same microglia, recorded on two consecutive days. Right panels: Zoom of one microglia at 0, 10 min and in a color-coded overlay with red (lost), green (gained), and yellow (stable) microglia processes. Lower panels (Zoom from upper panel): A lost microglial fine process is marked with a red arrow, a gained with a green arrow at 0 min and 10 min, respectively. **(D)** Microglial process motility depicted as fraction of gained, lost and stable microglial processes over a period of 30 min (n = 10). **(E)** Fractions of microglial fine processes on two consecutive days. One-way ANOVA with Šídák’s multiple comparisons test. F (2.331, 20.98) = 80.46; adjusted p-values: d0stable vs. d1stable=0.19; d0gain vs. d1gain=0.99; d0loss vs. d1loss=0.1, from n=10 individual microglia. **(F)** Turnoverrate of microglial fine processes.paired t-test, two-tailed, p=0.069, n=10 individual microglia **(G)** Coronal section 1.8 mm anterior bregma of the previously 3P in vivo imaged ROI with tdTomato expression in microglia and immunohistochemical staining for heat shock protein (HSP) and DAPI. **(H)** Coronal section 1.6 mm anterior bregma through the tumor from a glioblastoma mouse model with immunohistochemical staining for HSP, Microglia (Iba1) and DAPI. **(I)** HSP positive cells and microglia as percentage of DAPI^+^ cells per area in the 3P imaged ROI (ipsi.), contralateral hemisphere (cont.) and Glioblastom (GB). Avg. of 4 ROIs each ±std (ipsi.=1381 DAPI^+^ cells, cont.=1199 DAPI^+^ cells, GB=1248 DAPI^+^ cells).

### 3P-imaging of astrocytic Ca^2+^-activity in mPFC in vivo

Astrocytes are highly active during different brain states, with patterns ranging from spontaneous Ca^2+^-events in discrete microdomains to larger somatic Ca^2+^-changes and coherent astroglial network activity changes (*50, 51*). Cortical astroglial activity has so far been mostly recorded in superficial cortical layers, although several studies have indicated that the molecular and cellular repertoire of astrocytes strongly differs between superficial and deep cortical layers as well as different brain regions (*52, 53*). Therefore, we aimed to image astroglial Ca^2+^-activity in deep mPFC layers. We used the mouse line *GLAST-CreERT2::GCaMP5g::tdTomato* that conditionally (Tamoxifen-induced) expresses the Ca^2+^-indicator GCaMP5g and the reporter protein tdTomato in astrocytes in a Cre-dependent manner after tamoxifen induction (*54, 55*). We imaged mPFC through a cortical cranial window in head-fixed, anesthetized, 4-month old mice (**Fig. 4A**). Robust and specific td-Tomato reporter expression throughout all cortical layers was confirmed by post-hoc immunohistochemistry (**Fig. 4B**). We imaged astrocytes at 1300 nm excitation up to 1.2 mm deep from the brain surface, including the anterior cingulate and prelimbic areas (**Fig. 4C**; **movie S7)**. Ca^2+^-event analysis from deep cortical layers (1.0-1.2 mm from surface) showed that astrocytes in these layers displayed abundant spontaneous Ca^2+^ microdomains (**Fig. 4D-E**; **movie S8)**. The amplitudes, temporal kinetics and activity frequencies of these microdomains were similar to those reported for upper cortical layers in anesthetized mice using 2P-imaging (*56*). In addition, we also recorded Ca^2+^-changes from astrocytic somata (**Fig. 4F**). In general, these transients were larger, longer and less frequent than microdomains, consistent with 2P-imaging data from upper layer astrocytes in anesthetized mice (*57–59*). Hence, 3P-imaging enables recording of green-and red-fluorescent astrocytes at subcellular resolution in deep cortical layers. Moreover, our data suggest that, despite considerable layer-specific genetic heterogeneity, baseline Ca^2+^-activity in different cellular compartments is uniform across cortical layers in astrocytes.

**Fig. 4.**
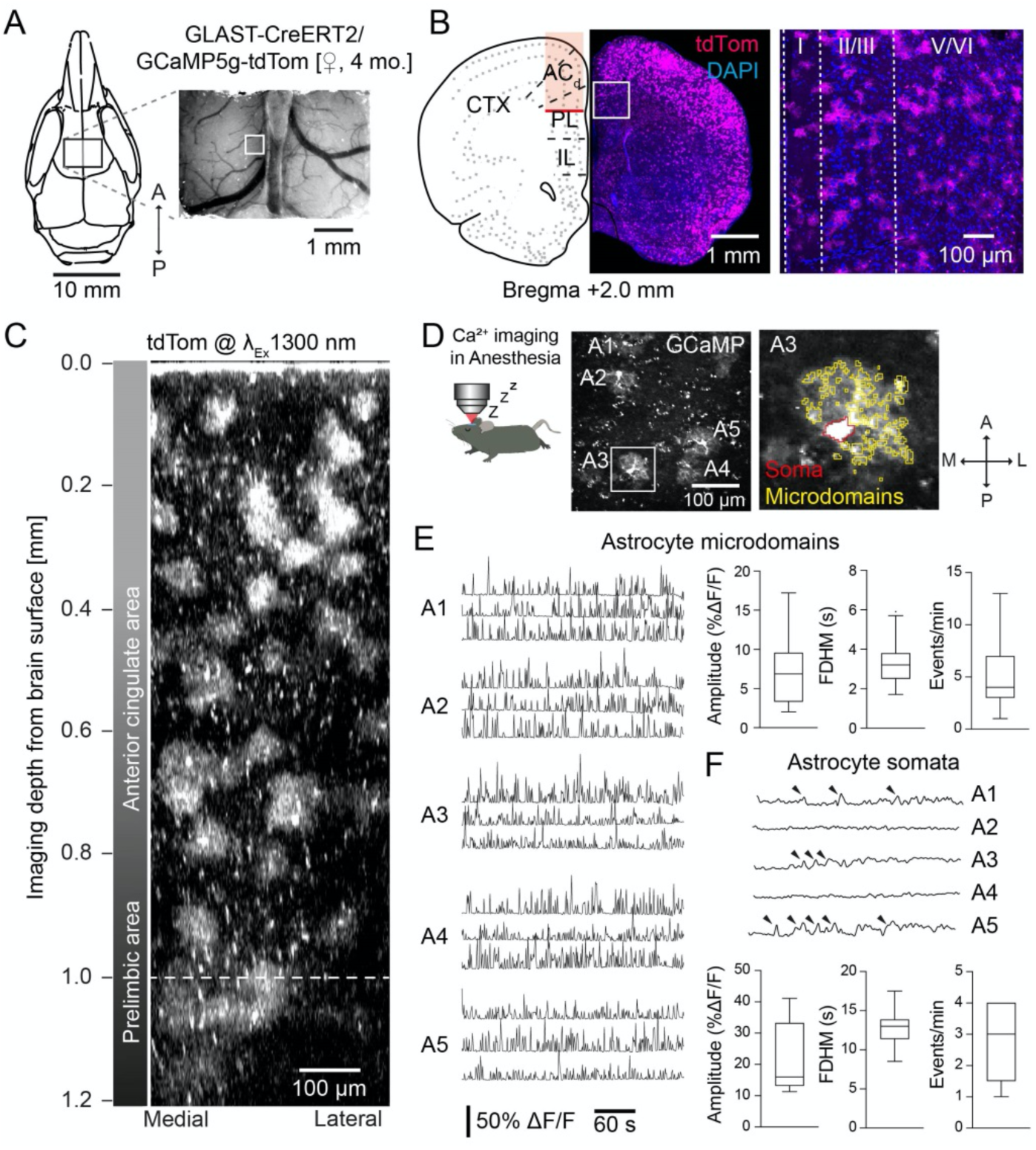
Astrocyte Ca2+-imaging in deep cortical layers. **(A)** mPFC imaging window position on the skull and representative image of the brain surface and vasculature (left, indicated ROI of z-stack shown in C); scheme of the anatomical localization of imaging window in relation to the mPFC subareas (right). **(B)** Coronal section 2 mm anterior to bregma, and expression pattern of tdTomato in astrocytes in mPFC. Magnification and cellular reporter expression in different layers of mPFC. **(C)** 3D reconstruction of 406 x-y frames from brain surface to 1200 µm below, acquired at a depth increment of 3 µm. **(D)** In vivo 3P imaging of GCaMP5g-positive astrocytes at 1000 µm below surface labeled A1-A5; left, zoom on A3 with automated ROI detection of the soma and active calcium domains. **(E)** Representative examples of calcium transients in astroglial microdomains recorded from A1-A5 with amplitude, full duration at half-maximum (FDHM) and event frequency (n = 10 astrocytes from n = 2 mice). **(F)** Calcium changes in astroglial somata (recorded from cells A1-A5; transients are labeled by arrowheads) with amplitude, FDHM and event frequency (n = 10 astrocytes from n = 2 mice).

### Imaging Ca^2+^-activity of neurons in the mPFC and dentate gyrus of awake head-fixed mice

Three-photon Ca²^+^-imaging of neurons has been performed in the cortex and hippocampus of mice, as well as in *Drosophila* (*26, 30*). In *Drosophila* we combined 3P-imaging with non-invasive mounting methods to carry out Ca^2+^-imaging in MB KCs in adult flies through the intact cuticle at cellular resolution confirming previous results **(Fig. S5, Suppl. Text 1)**. Ca^2+^-imaging in the mPFC of mice has been previously carried out through microprisms and GRIN-lenses (*39–41*). Here we employed three-photon Ca²^+^-imaging in the mPFC of awake head-fixed mice that expressed the Ca²^+^-indicator GCaMP6s in excitatory neurons (**Fig. 5A, B**). We used 1300 nm excitation to record somatic Ca^2+^-transients of VGluT2^+^ neurons in layer V of the prelimbic area of the mPFC as deep as 1.4 mm below the brain surface (**Fig. 5C**; **movie S9)**. Regions of interest (ROIs) were imaged with framerates ≥10Hz. We resolved and detected Ca^2+^-transients of more than 240 individual neurons in prelimbic layer V of the mPFC (**Fig. 5D, E**; **movie S10)**. The average Ca^2+^-transient amplitude was 4.95 ±4.14 [% ΔF/F] with a mean half width of 3.24 ±2.25 s (**Fig. 5F, G**). The spiking of each neuron was inferred from the corresponding Ca²^+^-traces (**Fig. 5E**) and a mean inferred spike frequency of 0.33 ±0.18 Hz was calculated for VGluT2^+^ neurons in prelimbic layer V of the mPFC in awake resting mice (**Fig. 5H**).

**Fig. 5.**
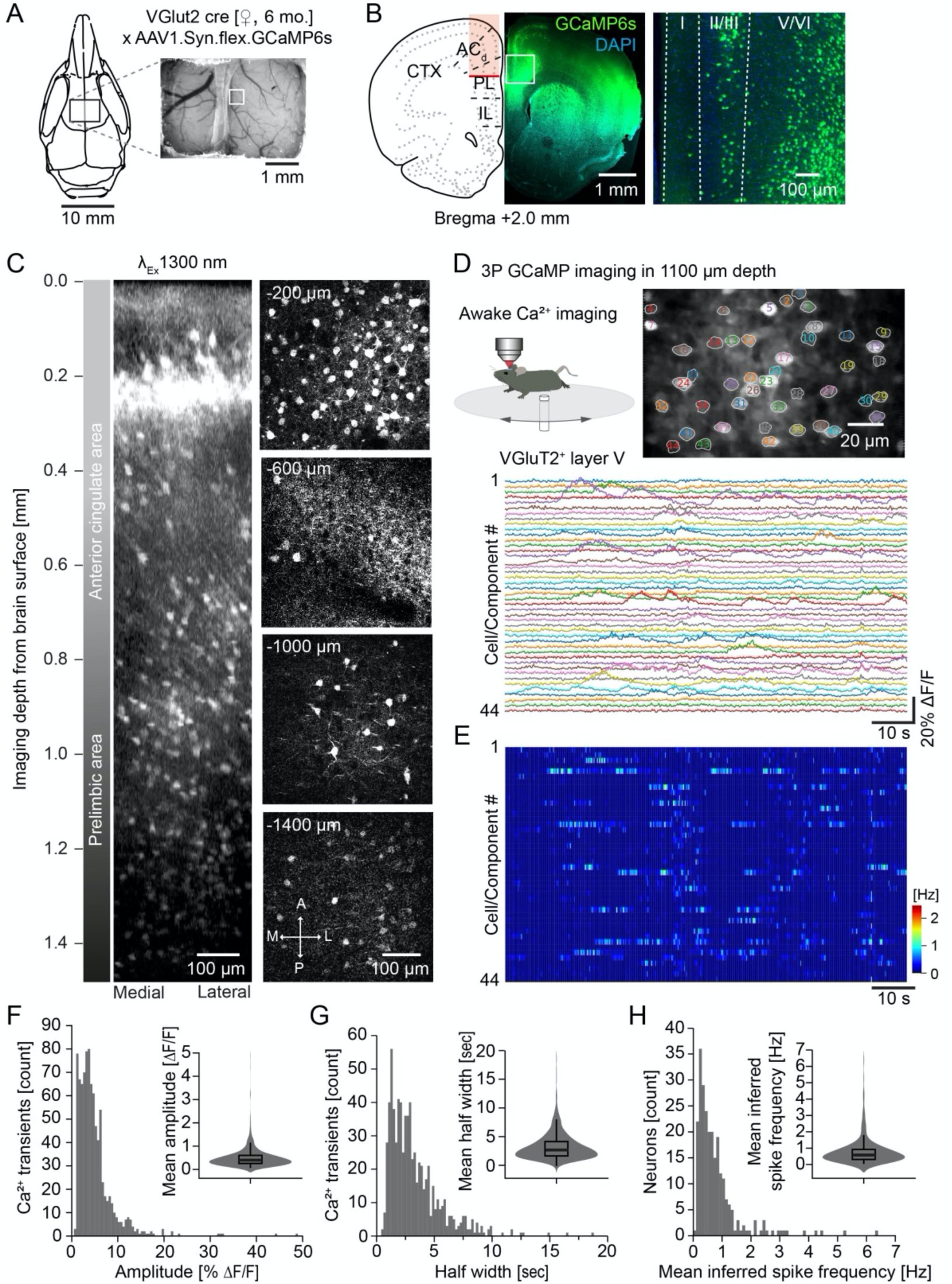
Awake Ca^2+^-imaging of excitatory neurons in the mPFC. **(A)** Schematic and exemplary picture of the cranial window placement in a vGlut2-Cre mouse to express GCaMP in glutamatergic neurons in the mPFC. **(B)** Schematic of a coronal section showing the imaged region in red (left). GCaMP6s expressing (green) neurons in the mPFC co-labeled with DAPI (blue). Overview (middle) and zoom (right) illustrating GCaMP6s expression in different cortical layers. **(C)** 3D reconstruction of a z-stack recorded from a depth up to 1420 µm below the brain surface (left). Exemplary images at various depths (right). **(D)** Schematic of 3P-imaging in an awake head-fixed mouse running on a circular disk (upper left). Average intensity projection with 43 ROIs of neurons in the mPFC at a depth of 1100 µm (upper right). Ca^2+^-recordings of the 43 labeled neurons (lower panel). **(E)** Inferred Ca^2+^-event rate (pseudo color coded) of the 43 neurons depicted in (D). **(F-H)** Number of Ca^2+^-transients over amplitude and mean Ca^2+^-transient amplitude (F). Number of Ca^2+^-transients over half width and mean half width (G). Number of neurons over mean inferred spike frequency and mean inferred spike frequency (H). (3 mice, 247 neurons, 882 Ca^2+^-transients)

Ca²^+^-imaging in the hippocampus of awake mice is a powerful tool to correlate neuronal activity with behavior. The dentate gyrus (DG) can be reached with 2P-imaging through an implanted hippocampal window **(Fig. S6A-C)**. Since SBR of 3P-imaging was superior beyond 500 µm depth (**Fig. 1G**), we performed 3P-imaging of DG neurons in transgenic GCaMP6f mice through an implanted hippocampal window **(Fig. S5D, E; movie S11)**. We imaged ROIs in DG with framerates ≥10Hz and precisely resolved sparse Ca^2+^-transients in head-fixed mice running on a linear treadmill **(Fig. S6E)**. These experiments underscore the applicability of 3P-imaging for awake Ca²^+^-imaging in deep brain areas like the DG and the mPFC, and provide the basic Ca^2+^-transient parameters of layer V excitatory neurons in the mPFC.

## Discussion

Here we applied three-photon *in vivo* imaging to record from different cell types and with different modalities in the prelimibic area of the mPFC. We demonstrated high-resolution sub-cellular imaging at depths below 1000 µm from the brain surface. Specifically, we recorded neurons at cellular resolution up to 1600 µm depth. We repetitively imaged dendritic spines of prelimbic layer 5 neurons over a week. We measured the processes’ motility of tdTomato-expressing microglia excited at 1650 nm in the prelimbic area of the mPFC. We carried out Ca^2+^-imaging in awake head-fixed mice to record Ca^2+^-transients from astrocytes and neurons in the prelimbic and infralimbic areas of the mPFC. Our data show, that three-photon *in vivo* imaging of prelimbic areas of the mPFC, without implantation of invasive GRIN lenses or microprisms, will be instrumental to elucidate the role of the mPFC in health and disease.

In the past, 3P *in vivo* imaging in rodents focused on reaching the hippocampus through the intact cortex (*25*). Young mice (<2 months) only have a thin and weakly myelinated corpus callosum (*60*). Therefore, the hippocampus is easily accessible even for less bright indicators used for Ca^2+^-imaging (*26, 61*). However, with age axons in the corpus callosum become highly myelinated, which increases light scattering, decreases excitation efficiency, and prevents image acquisition. Therefore, in older mice (>2 months) access to the hippocampus can instead be reached only from more lateral positions, where the excitation light does not need to pass through the thick myelinated corpus callosum **(Fig. S2, 4)**. Therefore, the dorsal hippocampal CA1 area is difficult to access with 3P-imaging in aged mice. This is especially the case for neurons expressing less brightly fluorescent GCaMP. The prelimbic area of the medial prefrontal cortex is positioned at a similar depth as the hippocampus, approximately 1 mm below the cortical surface. However, there is no myelinated fiber bundle separating the prelimbic area of the mPFC from above lying cortical areas. Indeed, we were able to visualize neurons 1700 µm deep in the mPFC, through a cranial window, without implanting GRIN lenses or microprisms previously used to access mPFC (*43, 44, 62*). Furthermore, we recorded dendritic spines with a size of 1-2 µm at >1000 µm depth. The same dendrites and spines of layer 5 pyramidal neurons in the prelimbic area of the mPFC were repetitively imaged over 7 days. We were able to measure the spine density on basal dendrites of layer V neurons (0.41 ±0.07 µm^-1^), which represents a significant improvement compared with a prism-implantation approach that limits spine imaging to apical dendrites in layer I and are more invasive. In comparison to GRIN lens approaches that are fixed to one region of interest (stably implanted), our approach with a 4 x 3mm window offers a larger brain region to be investigated. We would like to stress that no adaptive optics for wavefront shaping were necessary to resolve these tiny structures more than a millimeter deep. Adaptive optics have been used to improve the point spread function for sub-cellular resolution imaging in the hippocampus (*28, 31, 63*) and might also be beneficial for spine resolution improvement in the mPFC. However, adaptive optics are another costly device in the beam-path that potentially broaden the pulse width, which needs proper correction and adjustment. Our approach shows that subcellular resolution of dendritic spines, more than a millimeter deep, is achievable without adaptive optics.

Microglia are innate immune cells in the brain and rapidly react to small disturbances in the brain parenchyma (*10, 13*). Staining of microglia with a synthetic dye (Dylight649) enabled 3P-imaging at >1000 µm in the juvenile mouse brain (*64*). Furthermore, GFP-expressing microglia could be visualized through the skull with the application of adaptive optics (*65*). Here, we show 3P-imaging at 1650 nm excitation of td-Tomato expressing microglia in adult mice up to a depth of 1200 µm in the prelimbic mPFC area, which might be important for future experiments in mouse models of aging and neurodegeneration. In addition, we provide kinetics of microglial fine processes at subcellular resolution >1000 µm deep below the brain surface for the first time. Consecutive imaging of the very same microglia on subsequent days revealed unchanged kinetics and unperturbed morphology. Heatshock protein stainings remained negative, suggesting sufficiently low excitation energy. The measured turnover rate (∼55%) was comparable to previous measurements at smaller depths in cortex and hippocampus (*49, 66–68*), indicating that our imaging approach is well suited for long-term recordings of these sensitive cells.

Similar to microglia, astrocytes are glial cells of the brain that are highly sensitive to disturbances such as phototoxicity (*69*). Astrocytic Ca^2+^-imaging in anesthetized mice has been previously achieved in the corpus callosum applying 3P-imaging in combination with adaptive optics (*28*). Here we perform astrocytic Ca^2+^-imaging in the medial prefrontal cortex at >1000 µm depth without adaptive optics. We are able to resolve somatic signals as well as astrocytic calcium microdomains in the prelimbic area of the mPFC. Interestingly, although several studies have shown that the genetic composition of astrocytes differs strongly between superficial and deep cortical layers (*52, 53*), our data indicate that astroglial Ca^2+^-activity, at least under anesthetized conditions, is uniform across all cortical layers.

Awake Ca^2+^-imaging in mice, correlating neuronal network activity with behavior, has become a major tool to investigate brain function. In the hippocampus Ca^2+^-imaging and structural imaging has been established using implantation of tubes or GRIN lenses (*11, 12, 70*). With 2P-excitation, recordings from the dentate gyrus and CA3 region of the hippocampus are possible (*71–73*). Three-photon Ca^2+^-imaging in the dentate gyrus improved the image quality compared to 2P-imaging **(Fig. S6)**. However, it should be noted that due to the slower pulse repetition rates of 3P lasers (2-4 MHz) in comparison with 2P lasers (∼80 MHz), a smaller number of neurons can be imaged at 5-10Hz frame rates in the case of three-photon excitation (**Fig. 5**). Ca^2+^-imaging in the mPFC of awake mice has been enabled in the past by implantation of microprisms (*43, 44, 62*) or GRIN lenses in combination with two-and one-photon excitation (*62*). However, implantation of optics is usually at the expense of brain integrity. The implantation of a microprism limits access to superficial layers (1–3) of the medial prefrontal cortex. In contrast our approach is less invasive and gives access to deeper mPFC layers (3–5) that were previously out of reach. Therefore, 3P Ca^2+^-imaging in deep prelimbic and infralimbic areas of the mPFC will greatly advance our knowledge about neuronal ensembles and their relation to cognition in the future.

Here, we demonstrate sub-cellular three-photon *in vivo* imaging of dendritic spines, neurons, microglia and astrocytes in the prelimibic area of the mPFC of awake and anesthetized mice. We show that depth penetration to measure structural and functional dynamics of neurons and glial cells can be significantly improved by the application of 3P-imaging. As 2P-imaging in the past, 3P-imaging holds a great potential to revolutionize our understanding of the brain and especially the mPFC in the future.

## Materials and Methods

### Microscope

We have built a 3P setup as shown in (**Fig. 1A**). Three-photon excitation is achieved with a wavelength-tunable excitation source from Spectra-Physics (NOPA, Spectra-Physics) pumped by a femtosecond laser (Spirit, Spectra-Physics). Dispersion compensation has been realized with a prism-based compensator from APE (FemtoControl, APE). More details on the group delay dispersion of the setup have been published elsewhere (*74*). The laser repetition rate is maintained at 2 kHz. Laser intensity control was implemented by a lambda/2 plate (BCM-PA-3P, Thorlabs). A 50% underfilling of the objective’s back aperture was achieved with lens pairs (f:+100 and f:-50; AC254-100-C-ML and LD1464-C-ML, Thorlabs). The 2P excitation source is a Ti:sapphire laser (Chameleon UltraII, Coherent). The laser repetition rate is at 80 MHz. Images are taken with a multiphoton microscope built of a multiphoton microscope base (EMB100/M, Thorlabs), a motorized stage module (MCM3000, Thorlabs), and an epi-fluorescence LED illumination beam path (LED: MCWHLP1 and trinocular with 10x eyepiece: MCWHLP1, both Thorlabs). Primary scan path optics for Bergamo microscope series with the 900-1900 nm coating are used. Two GaAsP photomultiplier tubes (PMT2100, Thorlabs) in the 14° detector module (BDM3214-3P, Thorlabs) are used for non-descanned detection. The ThorDaQ with 3P Mezzanine Card (TDQ3-ESP, Thorlabs) is processing the data from the photomultiplier tubes and realizes the synchronization with the excitation laser pulses from the NOPA-Spirit laser with single pulse precision. ThorImage (Version 4.3, Thorlabs) is used to control imaging parameters. A 25x water immersion microscope objective with the numerical aperture of 1.05 (XLPLN25XWMP2, Olympus) is used. Green and red signals are separated by a 488 nm dichroic mirror (Di02-R488, Semrock) and 562 nm dichroic mirror (FF562-Di03). Then the GFP and third harmonic generated (THG) signals are further filtered by a 525/50 nm band-pass filter (FF03-525/ 50, Semrock) and 447/60 nm (FF02-447/ 60, Semrock) band-pass filter, respectively. Lateral movements are done with the 2D stepper motor (PLS-XY, Thorlabs), z-stacks are done using the built-in z device or a piezo focus device (PFM450, Thorlabs).

### Mice

Mice were group-housed and separated by gender with a day/night cycle of 12 hr. Water and food was accessible *ad libitum*. All experiments were performed according to animal care guidelines and approved by the Landesamt für Natur, Umwelt und Verbraucherschutz of North Rhine-Westphalia (Germany) (#81-02.04.2019.A084; #81-02.04.2018.A239; #81-02.04.2020.A059; 84-02.04.2017.A098). B6.Cg-TgN(Thy1-YFP-H)2Jrs mice (#003782) and B6.129P-CX3CR1^tm1Litt^/J mice (#005582) were purchased from The Jackson Laboratories. Thy1-YFP::CX3CR1-GFP mice were derived from a heterozygous crossing of CX3CR1-GFP and Thy1-YFP-H mice. Heterozygous Cx3cr1-creER/Rosa26_tdTomato/GFP.M mice were derived from crossing of Tg(Thy1-EGFP)MJrs/J mice (#007788), B6.129P2(C)-*Cx3cr1^tm2.1(cre/ERT2)Jung^*/J mice (#020940) and B6.Cg-*Gt(ROSA)26Sor^tm14(CAG-tdTomato)Hze^*/J mice (#007914) purchased from The Jackson Laboratories. Vglut2-Cre *Slc17a6^tm2(cre)Lowl^*/J mice (#016963), C57BL/6J-Tg(Thy1-GCaMP6f)GP5.5Dkim/J mice (#024276), B6;129S6-*Polr2a^Tn(pb-CAG-GCaMP5g,-tdTomato)Tvrd^*/J mice (#024477) and Tg(Slc1a3-cre/ERT)1Nat/J mice (#012586) were purchased from The Jackson Laboratories.

### Tamoxifen injections

To induce Cre recombinase expression in microglial cells, adult Cx3Cr1-CreER mice were injected intraperitoneally (i.p.) with 0.1 mg/g body weight tamoxifen for five consecutive days. Tamoxifen was dissolved in miglyol (Miglyol 812 Hüls Neutralöl) and administered in a volume of 5 µl/g body weight. To induce expression of GCaMP5g-tdTomato in GLAST-Cre::GCaMP5g-tdTomato-loxP mice, mice were injected i.p. at 5 μl/g with an emulsion of 20 mg/ml tamoxifen (Sigma) in sunflower oil:ethanol mix (Sigma) for five consecutive days.

### Cortical window preparation

Cortical window surgery was performed four weeks before imaging. According to each animal protocol, mice were either anesthetized with isoflurane (induction, 3%; maintenance, 1-1.5% vol/vol; Virbac), or with an i.p. injection of ketamine/xylazine (0.13/0.01 mg/g body weight). Body temperature was maintained with a heating pad at 37°C. Mice received buprenorphine (0.1 mg/kg; subcutaneously (s.c.), Reckitt Benckiser), dexamethasone (0.2 mg/kg; s.c., Sigma) and cefotaxime (2 g/kg; s.c., Fisher Scientific) shortly before the surgery. After fixation in a stereotaxic frame, the skin was removed under sterile conditions and a craniotomy above the prefrontal cortex (3x4 mm) or above the right somatosensory cortex (4 mm diameter) was created with a dental drill. The dura was carefully removed. The brain surface was rinsed with sterile saline, and adequate #1 coverslips (4 mm diameter or 3x4 mm) were sealed into the craniotomy with dental cement. For head-fixation during *in vivo* imaging a headpost (Luigs & Neumann) was cemented adjacent to imaging window. For analgesia, buprenorphine (0.1 mg/kg s.c.) was injected three times daily and Metamizol was (200 mg/kg) was applied to the drinking water for 3 consecutive days.

### Viral injections

Mice were anesthetized with an i.p. injection of ketamine/xylazine (0.13/0.01 mg/g body weight). After fixation in a stereotaxic frame and skin incision (5 mm), placement of the injection was determined in relation to bregma and a 0.5 mm hole was drilled through the skull. Stereotactic coordinates were taken from Allen brain reference atlas version 1 (2008). For Ca^2+^ imaging of mPFC layer 5 excitatory neurons 1 μl pAAV.Syn.Flex.GCaMP6s.WPRE.SV40 (Addgene) was injected bilaterally into mPFC of VGlut2-cre mice (+0.9 mm anterior-posterior, ±0.05 mm lateral and -1.4 mm ventral with a rostral/caudal 30° angle) at 0.1 μl/min, using a UltraMicroPump, 34G cannula and Hamilton syringe (World Precision Instruments, Berlin, Germany). After surgery buprenorphine (0.05 mg/kg) was administered three times daily for 3 days. mPFC window surgery followed two weeks after AAV injection.

### In vivo imaging

Mice were anesthetized with isoflurane (1–1.5% vol/vol) and head-fixed under the microscope on a heating pad at 37°C. Awake *in vivo* imaging was performed with habituated head-fixed mice on a rotating disc or on a linear treadmill. Mice were trained to run head-fixed on the linear treadmill. Velocity was read out by a rotation sensor, and the mouse’s actual position was computed with the help of a reflection light barrier. In addition, mice received one automatic liquid reward per lap (2m) at a fixed location.

### Image acquisition

Deep overview *in vivo* z-stacks were recorded with depth increments of 1-10 µm, 0.2-0.65 µm/pixel resolution and 2-3 µs pixel dwell time. Z-stacks of dendritic spines on basal dendrites of mPFC LV/VI neurons were imaged in 900-1100 µm depth with 1 µm depth increments, 0.15 µm/pixel resolution and 2 µs pixel dwell time. For the measurement of microglial fine process motility, z-stacks of individual microglia were imaged with 2-3 µm depth increments, 0.16 µm/pixel resolution, 2 µs pixel dwell time and with 5-10 min time-intervals for a period of 30 minutes. Timelapse imaging of astrocytic Ca^2+^-activity in mPFC in vivo was performed at 3 Hz frame rates with 1 µm/pixel resolution and 2 µs pixel dwell time. Imaging the Ca^2+^-activity of layer 5 excitatory neurons in mPFC of awake mice was performed at ≥10 Hz frame rates with 1-2 µm/pixel resolution and 2 µs pixel dwell time. Recordings of the Ca^2+^-activity of dentate gyrus granule cells in the hippocampus of awake mice was performed at 5-10 Hz frame rates with 0.5-1.5 µm/pixel resolution and 2 µs pixel dwell time. Spinal cord in vivo z-stacks were recorded with depth increments of 3 µm, 0.25 µm/pixel resolution and 2-3 µs pixel dwell time.

### Histology

Mice were deeply anesthetized with an i.p. injection of ketamine/xylazine and transcardially perfused with saline followed by 4% paraformaldehyde (PFA). Brains were taken out and coronal 70 µm slices were cut. Subsequently, after permeabilization (0.5% Triton-X100, 1h), slices were incubated with a DNAJ4/HSP40 antibody (1:100, mouse serum, Santa Cruz Biotechnology, sc-100714) and an Iba1 antibody respectively in the glioma condition (1:1000, rabbit serum, Wako, 019-19741) in a blocking reagent (4% normal goat serum, 0.4% Triton 1%, and 4% BSA in PBS) over night at room temperature. After washing the samples three times with PBS, secondary antibodies were administered (Alexa Fluor 647, goat anti-mouse, A21235, 1:400 and Alexa Fluor 488, goat anti-rabbit, 1:400, A11008) in 5% normal goat serum/BSA and incubated for 2h at room temperature. During the last 15 minutes of incubation DAPI was added (5mg/ml, 1:10000). Afterward slices were washed three times with PBS, mounted with Dako Mounting Medium, and covered with a glass cover-slip.

### Confocal Imaging

For the visualization of the different expression pattern and anatomical orientation, coronal slices were imaged with an LSM800 microscope. DAPI-(EX: G 365, Dichroic: FT 395, EM: BP 445/50), GFP-(EX: BP 470/40, Dichroic: FT 495 EM: BP 525/50), YFP-(EX: BP 500/20, Dichroic: FT 515, EM: BP 535/30) and tdTomato-(EX 561/10, Dichroic: 573, EM: 600/50) filter-sets were used. Large overview images were generated by stitching average intensity projections of x/y/z tile-scans with depth increments of 5-10 µm and 0.2-1.0 µm/pixel.

### Data analysis

#### Calcium imaging of neurons

For the analysis of neural activity in the mPFC and DG, a combination of tools taken from the python toolbox for calcium data analysis CaImAn (*75*), the python toolbox for spike inference from calcium data Cascade (*76*) and Igor Pro as well as custom-written code in python were used. Motion correction of the imaging recording was performed using the CaImAn-implementation for rigid-body registration. Detection of cell bodies and source-separation was performed using the CaImAn algorithm based on constrained non-negative matrix factorization (*77*) and the raw calcium signal was extracted from the ROIs. For DG recording, the ROIs were selected manually. ΔF/F was calculated using following equation: (F – F_0_)/F_0_ where F_0_ is the minimum 8^th^ quantile of a rolling window of 200 frames. Depending on the sampling rate of the recording, for spike inference the cascade algorithm trained to either the datasets Global_EXC_6Hz_smoothing200ms, Global_EXC_7.5Hz_smoothing200ms or Global_EXC_10Hz_smoothing50ms was used. For peak detection and determination of amplitude and half-width of decay we used Taro tools (https://sites.google.com/site/tarotoolsregister/).

#### Calcium imaging of astrocytes

Calcium imaging data were stabilized using a custom-written Lucas-Kanade algorithm (*78*) in Matlab R2018a (MathWorks). Timelapse recordings were down sampled to 1 Hz framerates by 3x frame averaging. Regions of interests (ROIs) representing astroglial microdomains were defined by fluorescence changes over time in GCaMP5g-positive astrocytes using a custom-written macro in ImageJ 1.50i (modified from (*56*)). Time-lapse data for each ROI were normalized, smoothed, and peak candidates were detected with a hard threshold. Detection and classification of fluorescence peaks over time was performed with a custom-written algorithm in Python. Mean fluorescence data were first normalized by a robust z-score calculated per ROI over the whole time-lapse series. Normalized data were then smoothed with a Gaussian filter, and all maxima above the threshold were selected as peak candidates. Peak candidates were defined by their ROI and the timepoint of peak maximum. Peak amplitude and full duration at half-maximum (FDHM) were determined for each peak candidate. Each time-lapse series was plotted together with the respective video file for visual inspection and verification.

#### Dendritic Spines

To remove noise in dendritic spine image data, a median-filter with a kernel size of 3x3 pixels was applied. The image data were subsequently stabilized using an optical flow method based on an iterative Lucas-Kanade solver (*79, 80*) from the open-source Python image processing library Scikit-image (*81*). The measurement of dendritic spine density was conducted by experienced human analyzers, who counted individual spines in each image stack and normalized the spine counts to the corresponding dendrite lengths (see, e.g., (*19, 49*)).

#### Microglia

Microglial fine process motility was measured with a custom analysis pipeline written in Python. First, noise was removed by applying a median-filter with a kernel size of 3x3 pixels. The individual consecutive time lapses of one mouse were maximum intensity projected along the z-axis and rigidly registered on each other using subpixel image registration by cross-correlation (*78*) provided by the open-source Python image processing library Scikit-image (*81*). Subsequently, the registered z-projections were binarized by thresholding the pixels in each projection using Otsu’s method (*82*). The thresholding was preceded by contrast limited adaptive histogram equalization (CLAHE) (*83*), also provided by the Scikit-image library, in order to enhance the contrast and, thus, the thresholding in each z-projection. Small blobs of less than 100 pixels were removed from the binarized images. We then calculated the temporal variation *ΔB(t_i_*) of each binarized image by subtracting the binarized image *B(t_i+1_)* at time point *t_i+1_* from two times the binarized image *B(t_i_)* at time point *t_i_*: *ΔB(t_i_) = 2×B(t_i+1_)-B(t_i_)* for *i=0, 1, 2, …, N-1*, where *N* is the total number of all time lapse time points. Pixels in *ΔB(t_i_)*, that have the value 1, were categorized as stable pixels, whereas pixels with the value -1 were categorized as gained pixels, and pixels with the value 2 as lost pixels. The microglial fine process motility was assessed by calculating the turnover rate (TOR) as the ratio of the number of all gained pixels *N_g_(t_i_)* and all lost pixels *N_l_(t_i_)* divided by the sum of all pixels: *TOR(t_i_) = (N_g_(t_i_)+ N_l_(t_i_)) / (N_s_(t_i_) + N_g_(t_i_)+ N_l_(t_i_)),* where *N_s_(t_i_)* is the number of all stable pixels. The average turnover rate &barTOR was culated by averaging *TOR(t_i_)* over all *N-1* time lapse time points.

#### Signal to background ratio (SBR)

Signal to background ratio was calculated for each frame individually by manually placing and measuring brightest signal-and lowest background ROI.

#### Cell counting

Single plane images of immunohistochemical stained brainslices were threshold adjusted, binarized and underwent watershed-algorithm based separation. Analysis was performed with Fiji (analysis of particles). >1000 DAPI^+^ cells were counted in four ROIs for each area.

#### Data accessibility

The authors declare that the data supporting the findings of this study are available within the paper, the methods section, and Extended Data files. Raw data are available from the corresponding author upon reasonable request.

### Statistical Analysis

Quantifications of microglial motility and subsequent statistical analysis, and graph preparation were carried out using GraphPad Prism 9 (GraphPad Software Inc, La Jolla, CA, USA). To test for normal distribution of data, D’Agostino and Pearson omnibus normality test was used. Statistical significance for groups of two normally distributed data sets paired or unpaired two-tailed Student’s t-tests were applied. One-way ANOVA with Šídák’s multiple comparison test was performed on data sets larger than two, if normally distributed. If not indicated differently, data are represented as mean ± SD. Figures were prepared with Illustrator CS5 Version 5.1 (Adobe) and IgorPro.6.37.

## Acknowledgments

We would like to thank Christof Moehl for providing support on astrocytic Ca^2+^-imaging data analysis.

## Author contributions

FF and MF prepared figures and wrote the manuscript with input from all authors. FF, FN carried out surgeries and recorded the data. FF and FM established the setup together with SK and HF. MM recorded data and analyzed Ca^2+^-imaging data. SP, MM, EAG and MM carried out measurements. BS and EB carried out spinal cord surgeries and recordings, partly together with FF. They wrote parts of the manuscript together with FB. ASL, ICWC carried out *Drosophila* measurements, prepared a figure and wrote parts of the manuscript together with GT. NR performed astrocytic measurements together with FF and performed astroglial calcium analyses. NR and GCP prepared a figure and wrote parts of the manuscript. SL carried out autocorrelator experiments together with FM. GCP, GT, FB and MF planned the project and supervised the work.

## Supplementary Text

### In vivo 3P imaging of the spinal cord

Like the corpus callosum, the spinal cord contains many myelinated axons. 3P-imaging in combination with AO significantly improved the penetration depth in comparison to two-photon imaging (*31, 46, 84*). Our results confirm previous findings and provide evidence that 3P-imaging in the dorsal horn of the spinal cord can be used to acquire subcellular resolution images at 300 µm depth without AO. It should be noted as for hippocampal preparations, that coverslip placement needs to be adjusted in a way that only few myelinated axons directly cross the light path. Imaging through highly scattering layers of myelinated axons degrades the point spread function and can only be partially corrected with AO.

### In-vivo imaging of the Drosophila Mushroom body developmental assembly and activity in response to odors, through the intact cuticle

Live imaging of the *Drosophila* brain, combined with recently available whole brain connectomes is a powerful tool to understand the organizational logic of fundamental circuits (*85, 86*). In *Drosophila*, the mushroom body (MB) is responsible for olfactory associative learning. Its input circuit, the calyx, includes the Kenyon cells (KCs) receiving olfactory information from the projection neurons (PNs) (*87*).

In adult flies, odor objects are represented by the sparse odor-specific activation patterns of the KCs population (*88, 89*). The major obstacle to functionally image the MB with high spatial resolution is the light scattering caused by fat bodies and trachea underneath the cuticle, that are thus typically removed. However, this increases the risk of damaging the MB underneath. Only recently successful preparations of adult flies were established leaving the cuticle intact in combination with 3P-imaging (*30, 90*). While this improvement created a path to longer term imaging, those preparations required partial compression of the fly’s head to achieve sufficient depth penetration. It is unclear whether flies survive that procedure for more than 24 hr. Therefore, with the use of 3P, we have developed a reversible mounting procedure **(Fig. S6A)** that effectively exposes the cell bodies of KCs from the posterior side **(Fig. S6B)**. 3P-imaging at 1300 nm excitation wavelength enabled deeper penetration through the fat body, resolving the small closely-packed KCs cell bodies (3-5 µm in diameter) **(Fig. S6C)**. We performed Ca²^+^-imaging of KCs expressing GCaMP6f and a nuclear marker (NLS-Cherry) in male flies **(Fig. S6C).** Flies were head-fixed under the 3P-imaging setup and stimulated with individual odors **(Fig. S6D)**. Segmentation of KCs was carried out based on nuclear NLS-mCherry expression **(Fig. S6E, movie S13)**. Corresponding dF/F_0_ of the GCaMP signal was calculated for individual

KCs **(Fig. S6F)**. KCs that responded to the same odor for all three trials were considered as responding units **(Fig. S6G)**. Here, we provide a refined 3P-imaging approach that does not include head compression, while allowing resolution of individual mushroom body neurons and recording of their Ca^2+^-transients. This approach will be useful to correlate neuronal activity with long-term behavior experiments in *Drosophila*. Thereby, the toolkit to understand the fly brain will be greatly advanced.

## Supplemental Methods

### Spinal cord window implantation

A female 12-week-old GFP-M mouse was administered buprenorphine (Temgesic, s.c, 0.1 mg/kg) and anaesthetized with ketamine-xylazine (i.p, 100 mg and 10 mg/kg respectively). Fur was shaved and skin sterilized with sequential iodine and ethanol swabs before transferring the mouse to a self-regulated heating pad at 37°C. An incision was made over thoracic spinal cord. Muscle was retracted and a dorsal laminectomy of thoracic spinal level 12 was performed. Dura was removed using forceps (Dumont 5) and angled spring scissors (FST 15010-09). The vertebral column was stabilized using a multi-joint frame and adson forceps. A 3x4 mm coverslip was secured using UV-cured adhesive (Norland Products) and dental cement.

### Fly Stocks

Flies were raised in a 12h/12h light-dark cycle on a standard cornmeal-based diet at 25 °C, 60

% relative humidity. All functional imaging experiments were performed on adult male flies 3–5 days after eclosion. The genotype of flies for imaging KC activity was *+/+;UAS-GCaMP6f, UAS-nls-mCherry/+;VT030559-Gal4/+*. *VT030559-Gal4* is a driver for most of the KCs in the mushroom body. *UAS-GCaMP6f* encodes GCaMP6f while *UAS-nls-mCherry* encodes for mCherry with a nuclear localization signal, as reported in Delestro et al. (*91*).

### Ca^2+^ imaging of the Drosophila MB KCs with intact cuticle

Male adult flies were briefly anesthetized on ice, and then placed on a chilled aluminum block. The posterior part of the head capsule was positioned upward, and attached to the edge of a glass coverslip using UV-curable glue (NOA 68, Norland Products Inc.). As the KC cell bodies are closer to the cuticle on the posterior side compared to the dorsal side of the head, this mounting method could effectively expose the cell bodies, and reduce light scattering through a thinner layer of fat body. Then, the glass coverslip with the head-fixed fly was positioned carefully in and fixed to the imaging chamber. The flies were then imaged with the 3P microscope setup described here and equipped with a 25× Olympus XLPlan N WMP2 (1.05 NA, 2.0 mm WD) water-immersion objective. Both GCaMP6f and nls-mCherry were excited at 1300nm. Bandpass filters for 525/550 nm and 593/620 nm were used for detecting the GCaMP and mCherry fluorescence signals, respectively. Individual monomolecular odors were used to stimulate the olfactory responses of KCs. Odor stimuli were delivered to the fly using a 220A olfactometer (Aurora Scientific). The odors 3-Octanol (OCT) and 4-Methylcyclohexanol (MCH) were diluted at 1:100 and 1:10 in mineral oil, respectively. Upon delivery to the fly, the olfactometer further diluted the odor in air, resulting in the final dilution of 1:1000 for OCT and 1:100 for MCH. Three trials of 3 s odor pulses of OCT, MCH, or mineral oil were delivered to the animal in a randomized order, with 35 s constant flow of pure air between odor exposures. After each imaging session, the fly can be detached from the coverslip and return to the standard food vial with other flies.

### Image analysis of the Drosophila MB Ca^2+^-imaging

Image processing was conducted using Fiji, and data analysis was performed with custom codes written in python. For the MB calcium imaging, the 2D time series was first motion-corrected using the ‘Template matching and slice alignment’ plugin in Fiji, which performed slice registration with a selected landmark. Afterwards, the cell bodies were identified using the nls-mCherry signal and the segmentation function of the ‘Trackmate’ plugin in Fiji, followed by manual adjustment. To quantify the neural responses along the time series, we extracted the fluorescence intensity of individual KCs for both the nls-mCherry and the GCaMP6f channels. The baseline fluorescence level (F_0_) was calculated by averaging the 17 frames before the first odor delivery for each channel. dF/ F_0_ for both channels were then computed using (F_t_ -F_0_)/ F_0,_ respectively. The axial-brain-motion-induced fluorescence change at the GCaMP channel was corrected with the use of the dF/ F_0_ from the nls-mCherry channel. To be considered as responsive to a particular odor, the peak dF/ F_0_ within 10 frames after that odor release should be above the responding threshold (2STD). Only those KCs that respond to the same odors across all three trials are considered as responding units.

### Hippocampal window preparation

To image dentate gyrus granule neurons, a hippocampal window was implanted above the right dorsal hippocampus as described before (*12*). Anesthesia was established with an i.p. injection of ketamine/xylazine (0.13/0.01 mg/g body weight). Additionally, buprenorphine (0.05 mg/kg s.c.) was injected shortly before the surgery. For head-fixation during in-vivo imaging a headpost (Luigs & Neumann) was cemented adjacent to the hippocampal window. Analgesia was carried out with buprenorphine injections (0.05 mg/kg s.c.), three times daily for 3 consecutive days. In-vivo imaging started after 4 weeks of recovery.

**Fig. S1:**
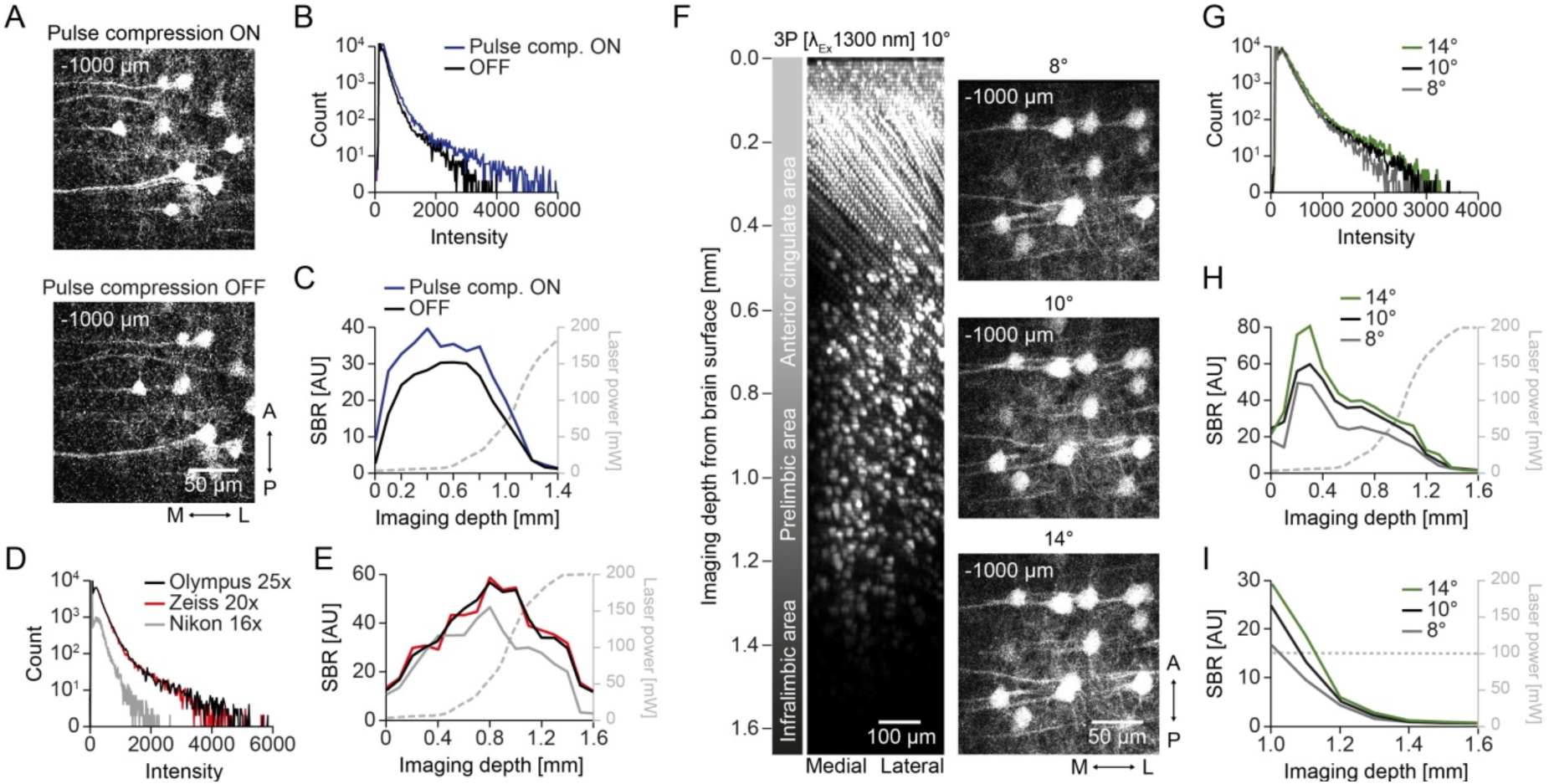
Optomechanical adjustments to improve the microscope. **(A)** Exemplary images of z-sections acquired at 1 mm depth with pulse compensation ON and OFF. **(B)** Comparison of fluorescence histograms at 1 mm depth with or without pulse compression. **(C)** SBR as a function of imaging depth for the z-stacks recorded with or without pulse compression including laser-power as a function of imaging depth. **(D)** Comparison of fluorescence histograms at 1 mm depth for the three different objectives (Olympus 25x, Zeiss 20x and Nikon 16x) **(E)** SBR as a function of imaging depth for the z-stacks recorded with the three different objectives including laser-power as a function of imaging depth. **(F)** 3D reconstruction of a z-stack recorded from a depth up to 1600 µm below the brain surface (left). Exemplary images at 1 mm depths (right) for 8°, 10° and 14° modification on the detection optics. **(G)** Comparison of fluorescence histograms at 1 mm depth for 8°, 10° and 14° modification on the detection optics. **(H)** SBR as a function of imaging depth for the identical z-stacks recorded with 8°, 10° and 14° modification on the detection optics, including laser-power usage as a function of imaging depth. **(I)** SBR as a function of imaging depth for the identical z-stacks recorded with 8°, 10° and 14° modification on the detection optics at constant laser-power (100 mW).

**Fig. S2:**
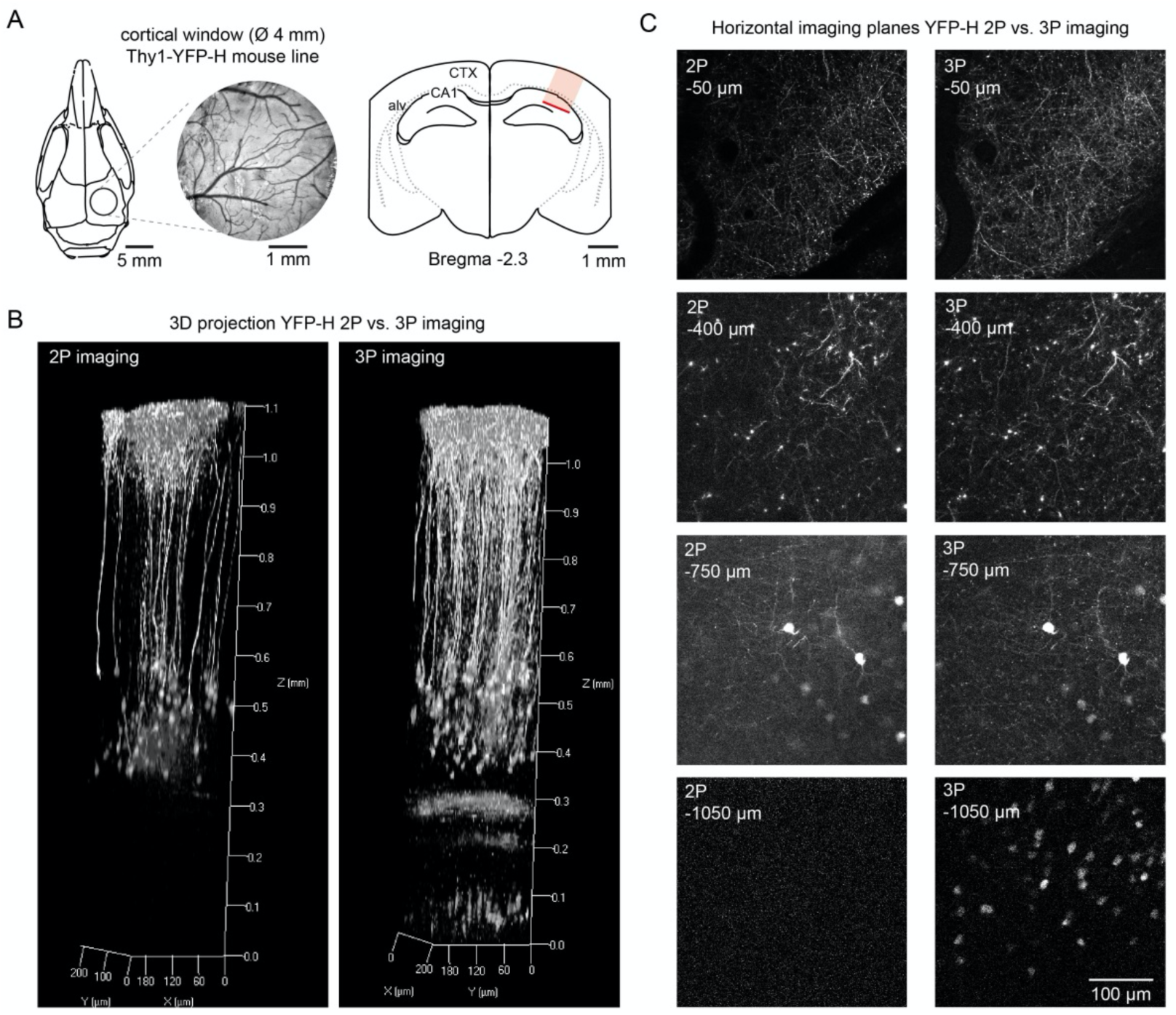
2P-versus 3P-imaging through the cortex into the hippocampus. **(A)** Schematic of the cranial window position above the somatosensory cortex (left), and red marked area of the imaging region on a coronal section (right). **(B)** 3D reconstruction of a z-stack acquired with 2P (left) versus 3P (right) imaging in a Thy1-YFP-H mouse (4 month old), expressing YFP in a subset of excitatory neurons. **(C)** Exemplary images of z-sections acquired at different depth comparing 2P with 3P excitation.

**Fig. S3:**
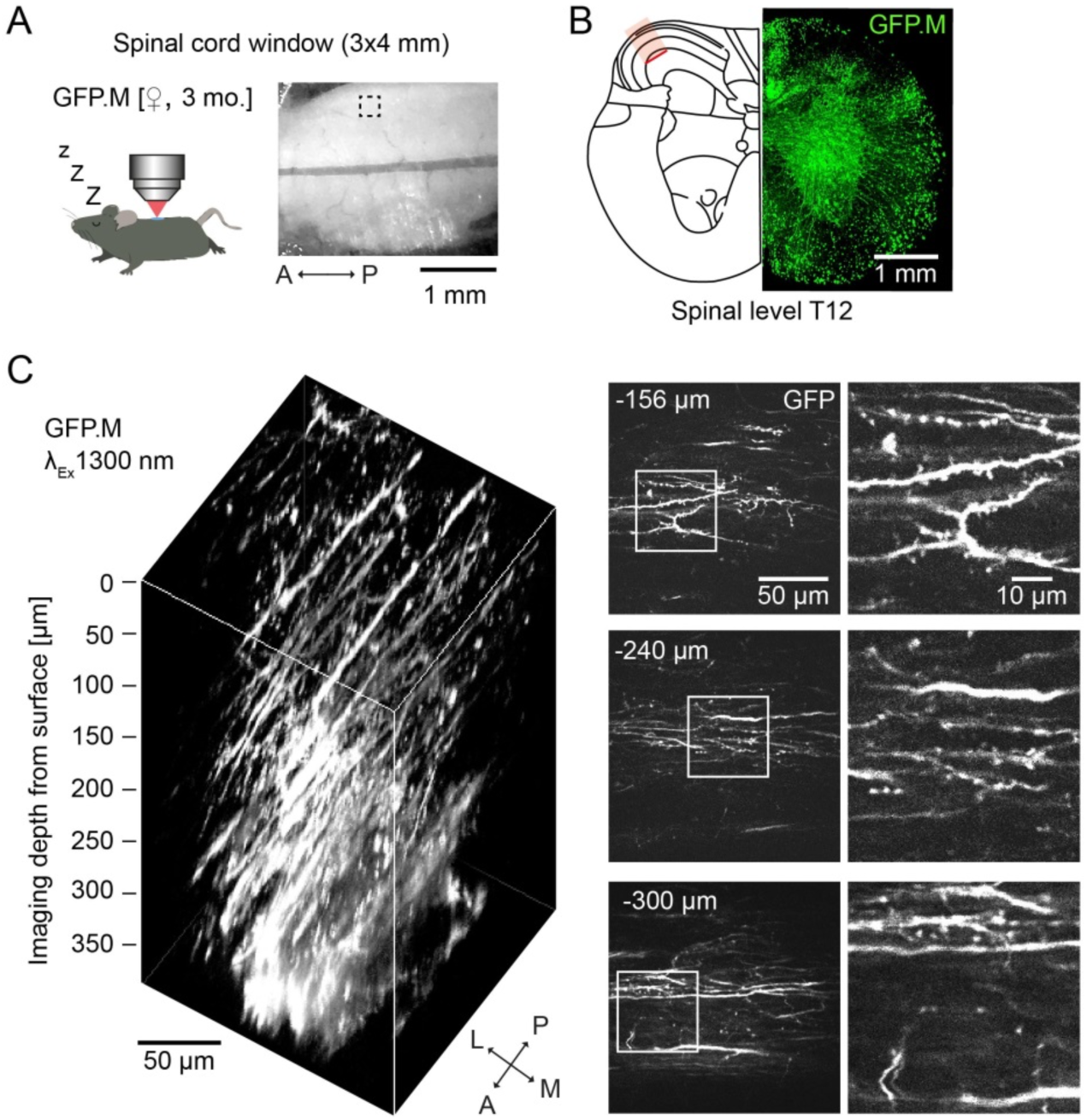
In vivo 3P imaging of the spinal cord. **(A)** Schematic showing spinal cord in vivo imaging in an anesthetized GFP-M mouse and a picture illustrating the view into the implanted spinal cord window with indicated imaging ROI position. **(B)** Schematic of a transverse section spinal level T12 (left panel) and a confocal microscopy picture from a corresponding section in a GFP.M transgenic mouse (right panel). **(C)** 3D reconstruction of 132 x-y frames from spinal cord surface to 390 µm below taken at a depth increment of 3 µm with 1300 nm Excitation (left) and exemplary individual z-planes recorded at different depths as indicated in the pictures with magnified ROIs (right).

**Fig. S4:**
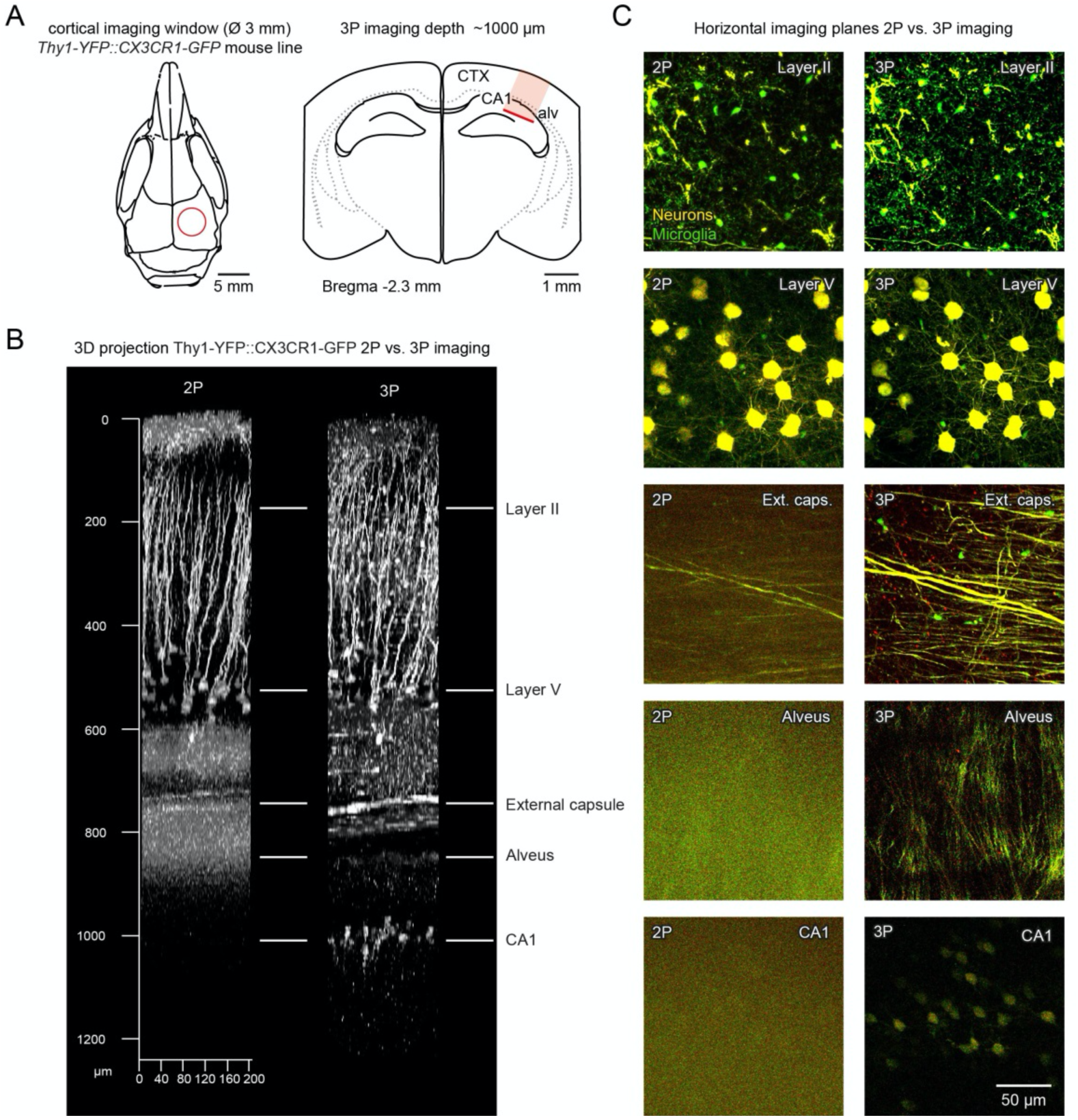
2P-versus 3P-imaging through the cortex into the hippocampus. **(A)** Schematic of the cranial window position above the somatosensory cortex (left), and red marked area of the imaging region on a coronal section (right). **(B)** 3D reconstruction of a z-stack acquired with 2P (left) versus 3P (right) imaging in a Thy1-YFP-H::Cx3cr1^GFP^ mouse (4 month old). **(C)** Exemplary images of z-sections acquired at different depth comparing 2P with 3P excitation. Microglia are labeled in green (GFP) and neurons are labeled in yellow (YFP).

**Fig. S5:**
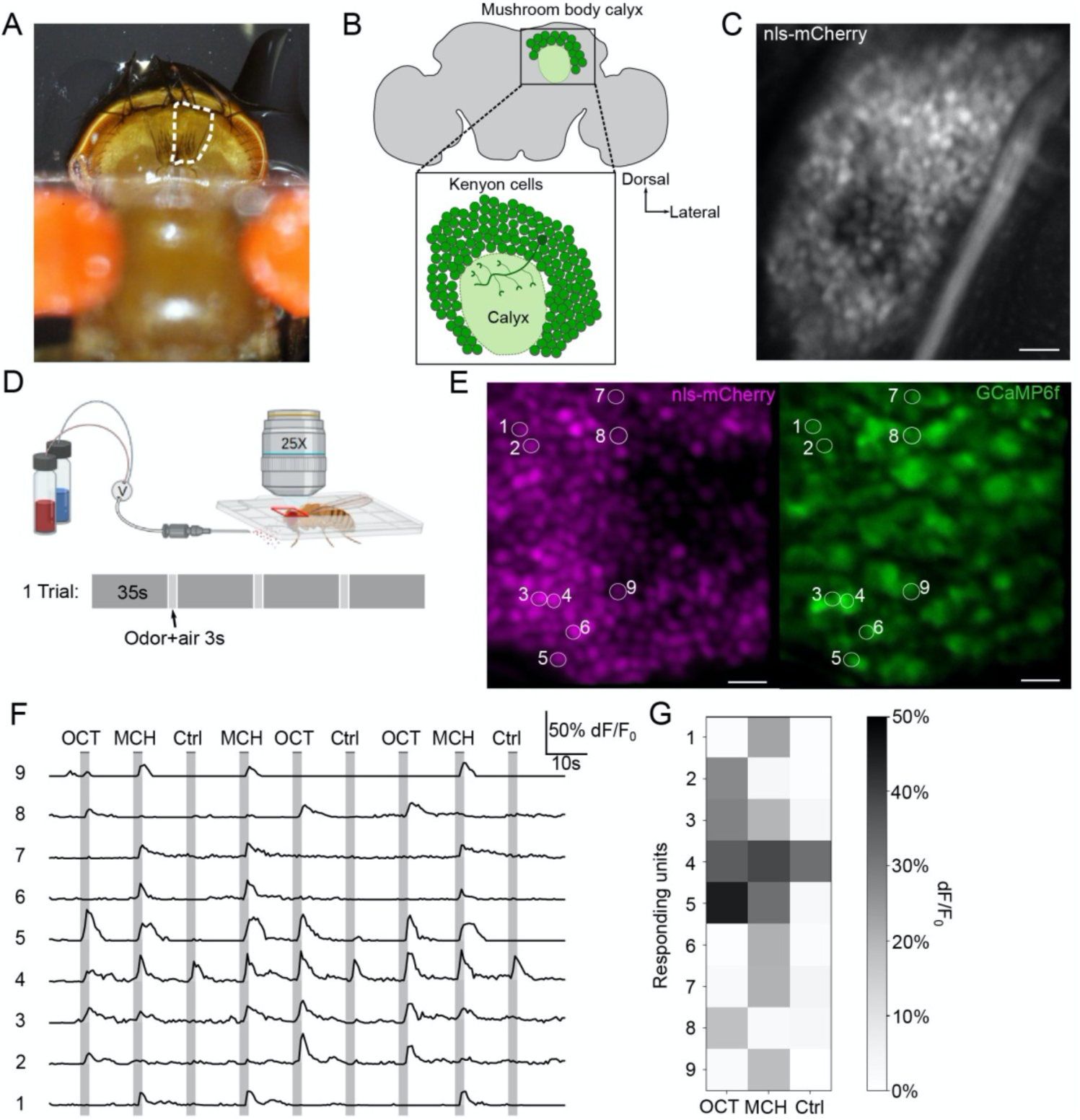
*In-vivo* functional imaging of the *Drosophila* Mushroom body with intact cuticle. **(A)** Image of a fly attached to a glass coverslip, exposing the posterior part of the head capsule for functional imaging of the KCs (the dotted line region). **(B)** Schematics of the adult fly brain, with the highlighted MB calyx. The KCs cell bodies (green circles) surround the MB calyx, where their dendrites receive PN olfactory inputs. The imaging plane (insert) contains around 150 KCs cell bodies across the calyx. **(C)** Example volume of the KCs nuclei expressing nls-mCherry (20μm thick with 40 sections). Scale bar, 10μm. **(D)** Scheme of the experimental setup and odor stimulation protocol (adapted from Prisco et al., 2021). 3 s of odor delivery of each of the 3 odors (3-Octanol (OCT), 4-Methylcyclohexanol (MCH) and mineral oil (Ctrl)), inter-odor interval of 35 s. **(E)** Left panel: KCs nuclei labelled with nls-mCherry. Right panel: Cell bodies of KCs expressing GCaMP6f. The activity of each circled responding unit is shown in (F). Scale bar, 10 µm. **(F)** Time course of response (dF/F0) of the responding units to a series of odor stimuli. The numbers correspond to the circles highlighted in (E). **(G)** Trial average peak response of the responding units.

---

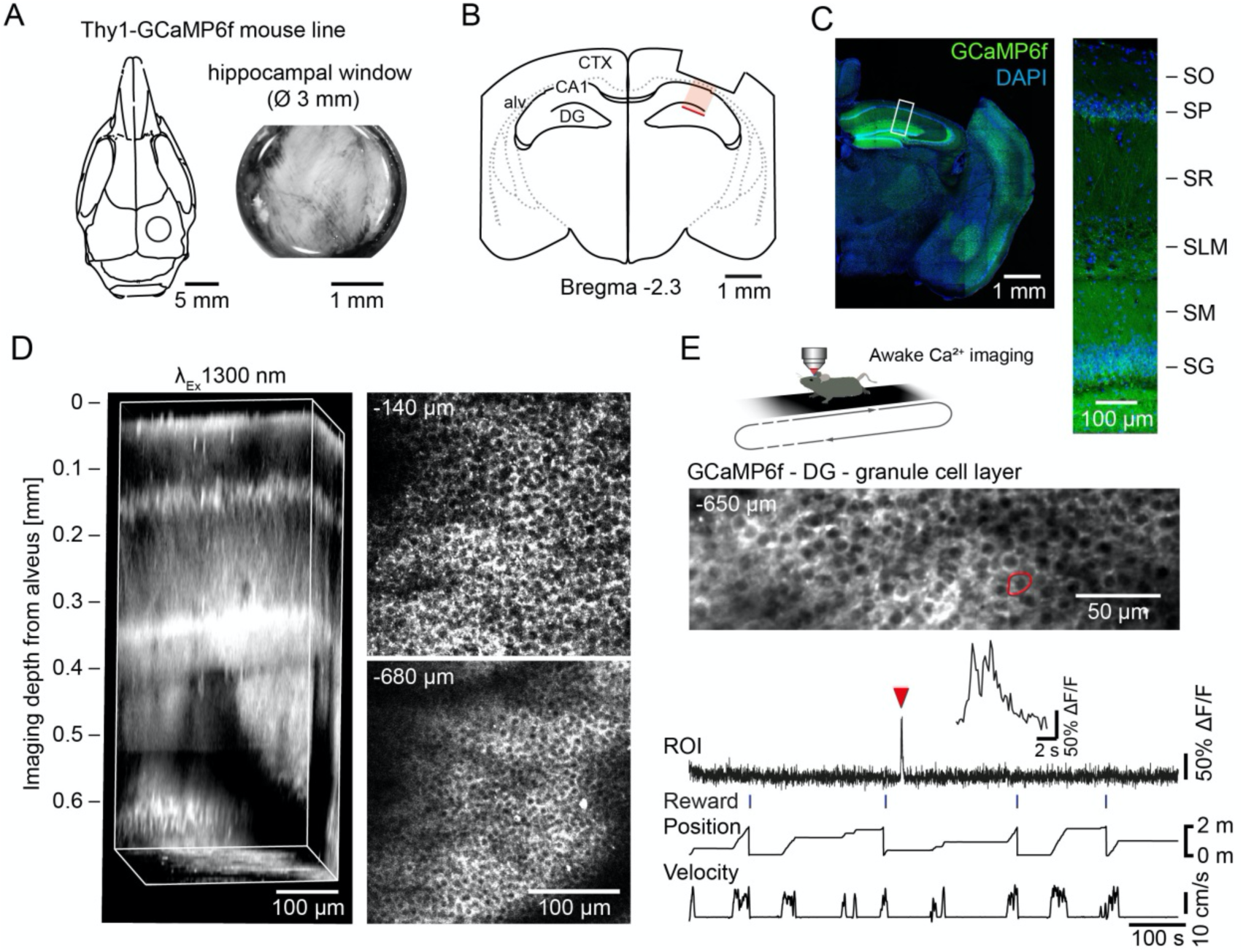

**Movie S1.**

In-vivo 3P z-scan of 320 x-y frames from brain surface to 1600µm below, taken at a depth increment of 5µm in the mPFC of YFP-H transgenic mouse.

**Movie S2.**

In-vivo 2P mPFC z-scan with 920nm excitation in a YFP-H transgenic mouse.

**Movie S3.**

In-vivo 3P mPFC z-scan with 1300nm excitation in a YFP-H transgenic mouse.

**Movie S4.**

In-vivo 3P mPFC z-scan with 1300nm excitation in a Thy1-GFP-M transgenic mouse.

**Movie S5.**

In-vivo 3P mPFC z-scan with 1650nm excitation in a Cx3Cr1-creER2 Rosa tdTomato mouse on day0.

**Movie S6.**

In-vivo 3P mPFC z-scan with 1650nm excitation in a Cx3Cr1-creER2 Rosa tdTomato mouse on day1.

**Movie S7.**

In-vivo 3P z-scan of 406 x-y frames from brain surface to 1200 µm below, acquired at a depth increment of 3 µm in the mPFC of a GLAST-CreERT2::GCaMP5g::tdTomato transgenic mouse.

**Movie S8.**

In vivo 3P recording of GCaMP5g-positive astrocytes at 1000 µm below surface.

**Movie S9.**

In-vivo 3P z-scan from brain surface to 1420µm below in a vGlut2-Cre mouse expressing GCaMP6s in glutamatergic neurons in the mPFC.

**Movie S10.**

In-vivo 3P recording of GCaMP6s-positive glutamatergic neurons in the mPFC at a depth of 1100 µm.

**Movie S11.**

In-vivo 3P z-scan from SO to SG up to 700 µm deep into the dorsal hippocampus through a hippocampal window in a Thy1-GCaMP6f transgenic mouse.

**Movie S12.**

In-vivo 3P z-scan of 132 x-y frames from spinal cord surface to 390 µm below taken at a depth increment of 3 µm with 1300 nm excitation in a Thy1-GFP-M transgenic mouse.

**Movie S13.**

In-vivo 3P functional imaging of the *Drosophila* Mushroom body with intact cuticle.

